# Curvilinear features are important for animate/inanimate categorization in macaques

**DOI:** 10.1101/2020.08.25.267393

**Authors:** Marissa Yetter, Sophia Robert, Grace Mammarella, Barry Richmond, Mark A. G. Eldridge, Leslie G. Ungerleider, Xiaomin Yue

## Abstract

The current experiment investigated the extent to which perceptual categorization of animacy, i.e. the ability to discriminate animate and inanimate objects, is facilitated by image-based features that distinguish the two object categories. We show that, with nominal training, naïve macaques could classify a trial-unique set of 1000 novel images with high accuracy. To test whether image-based features that naturally differ between animate and inanimate objects, such as curvilinear and rectilinear information, contribute to the monkeys’ accuracy, we created synthetic images using an algorithm that distorted the global shape of the original animate/inanimate images while maintaining their intermediate features (Portilla and Simoncelli, 2000). Performance on the synthesized images was significantly above chance and was predicted by the amount of curvilinear information in the images. Our results demonstrate that, without training, macaques can use an intermediate image feature, curvilinearity, to facilitate their categorization of animate and inanimate objects.

## Introduction

Primates can recognize objects with remarkable speed and accuracy—an ability that is crucial for avoiding predators, identifying food sources, and otherwise surviving in their natural habitat. Though seemingly effortless, decades of research in visual neuroscience and computer vision have shown that the ability to extract an object from a visual scene and categorize it is far from trivial (e.g. Pinto et al., 2008). The primate brain is equipped to deal with this computational problem by exploiting a vast array of features to classify objects into categories. Some distinctions are made based on knowledge or experience with the object, such as how it can be used (Bovet & Vauclair, 1998; Träuble & Pauen, 2007), whether it is threatening (Lipp, 2006; LoBue & DeLoache, 2011), or what contexts it is often found in (Kalénine et al., 2009, 2014; Blake et al., 2007), while others are determined based on the appearance of the object alone, by using its visual features such as color, size, global shape, and texture, etc.

The relative contribution of knowledge- and image-based information to object categorization varies across situations due to a number of factors. A crucial factor is the extent to which image-based features are predictive of a meaningful category or object class—a reasonable prerequisite for a visual system to rely on visual cues for object classification. Furthermore, the category or object class itself might influence the relative contribution of image information and prior experience needed to perform categorization. A long-standing line of research in evolutionary psychology has suggested that the primate visual system is highly tuned for the detection and recognition of animacy (Nairne et al., 2017; Meyerhoff et al., 2014; Calvillo et al., 2016; Long et al., 2019), even as early as 3 months old (Heron-Delaney et al., 2011; Opfer & Gelman, 2011; Rakison, 2003). A number of biological processes and key image feature differences have been proposed to explain how this discriminative ability might emerge so early in development. For example, some researchers have argued that innate processing biases interact with crude image-based biological templates to produce a sensitivity to faces from birth (Chiara et al., 2008; Sugita, 2008). Others have argued for a greater emphasis on the role of experience, through which persistent social exposure to faces early in life leads to a preference for face stimuli via more domain-general neural mechanisms (Livingstone et al., 2017; Srihasam et al., 2014). Yet another line of research has shown that human infants might develop concepts of animacy based on differences between biological and non-biological motion (Simion et al. 2008; Mandler, 1992).

That the animate-inanimate distinction might be special to our visual system, and that these two categories differentially covary with a number of image features, suggests a plausible mechanism by which the primate visual system evolved to exploit image feature covariances, such as those listed above, to make animate-inanimate categorization judgments. One such feature is curvilinearity, or the extent to which the image of an object is composed of curved lines and textures. Animate objects tend to be more curvilinear than inanimate objects (Kurbat, 1997; Levin et al., 2001). A recent study by Zachariou et al. (2018) demonstrated that, when deprived of global shape cues, humans were able to categorize animate and inanimate objects using just curvilinear information. Further, curvilinear information was positively correlated with performance on images of animate objects and negatively correlated with performance on inanimate objects. Given the lack of object shape information in the stimuli used and the lack of relationship between subjects’ confidence ratings and their accuracy, it appears that this categorization ability is driven by an implicit, primarily bottom-up process.

If the human visual system can implicitly rely on curvilinear information to perform animate-inanimate categorization, it is possible that this may be a property of the primate visual system more broadly. To test this hypothesis, the current study sought to establish the contribution of image-based information to animate-inanimate categorization in a non-human primate, the rhesus macaque, by: (1) testing the ability of macaques to categorize a large trial-unique set of animate and inanimate intact images that were unfamiliar to them; and (2) testing whether the macaques could use curvilinearity, without training, to categorize the objects when global shape information was removed.

## Materials and Methods

### Subjects

Three male rhesus macaques (5 - 8 kg) were used in two behavioral experiments. All experimental procedures were approved by the National Institute of Mental Health Animal Care and Use Committee.

### Visual stimuli

The first experiment included 500 images of animate objects and 500 images of inanimate objects which were downloaded from open-source repositories on the internet. The animate images were comprised of mammals, birds, fish, reptiles, and insects (Figure 1a). The inanimate images included human-made objects such as tools, vehicles, buildings, various household items, and natural objects, such as rocks and flowers (Figure 1b). All object images were digitally processed (see Supplementary Materials for a detailed description of this process) to match size, background, mean luminance and root-mean-square (RMS) contrast. All images were resized to 200 × 200 pixels.

**Figure 1:**
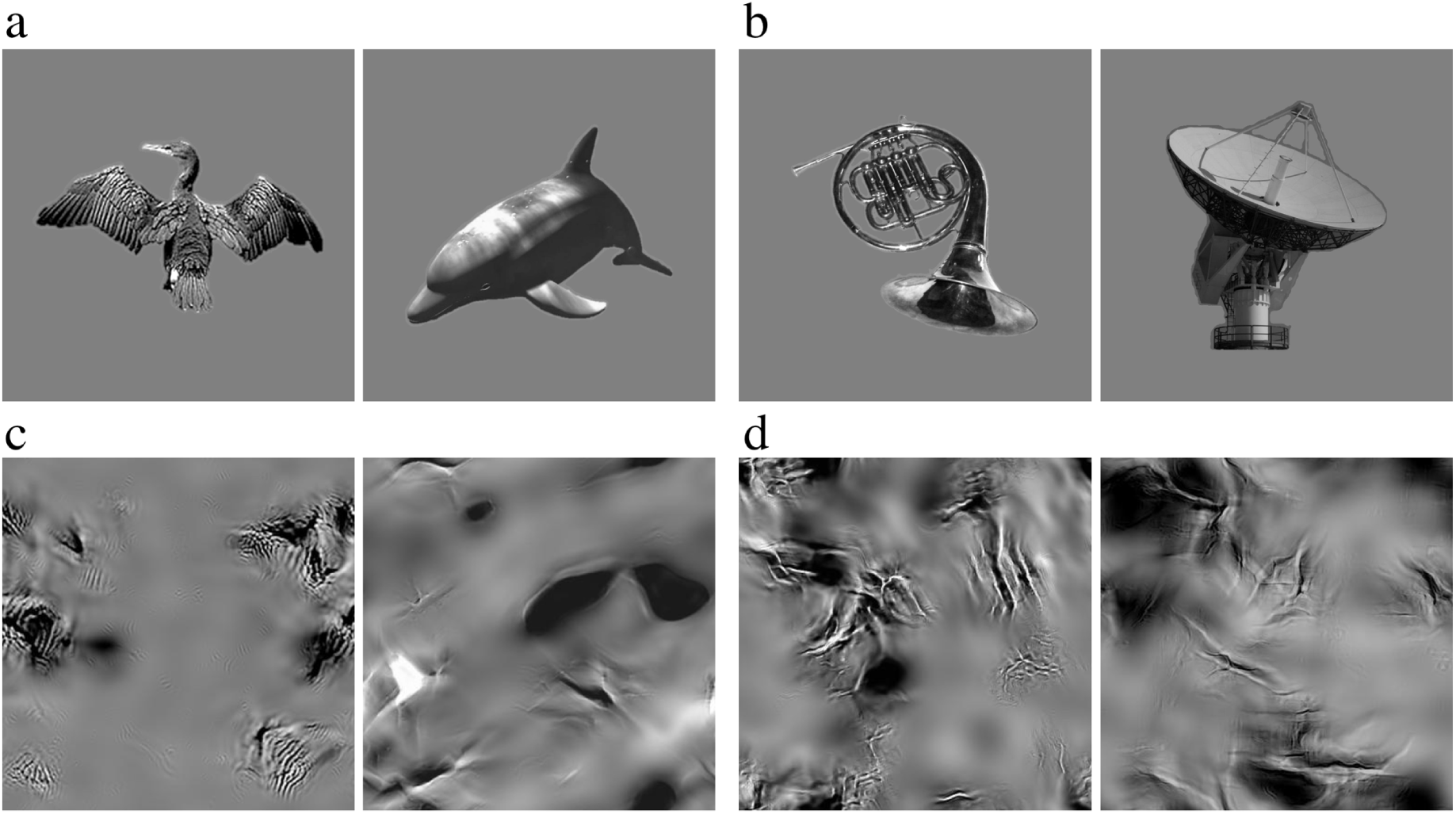
Examples of stimuli: (a) animate images; (b) inanimate images; (c) synthesized animate images; (d) synthesized inanimate images.

For the second experiment, we used an algorithm, described in detail in Portilla and Simoncelli (2000), to generate synthesized images of animate and inanimate objects (Figure 1c and 1d) that abolished the global shape of the original images but maintained their intermediate visual features (see Supplementary Materials). 1000 synthesized images were generated using the testing set of 500 animate and 500 inanimate intact images used in Experiment 1.

### Experimental procedures

The monkeys sat in a primate chair inside a darkened, sound-attenuated testing chamber. They were positioned 57 cm from a computer monitor (Samsung 2233RZ, Wang and Nikolic 2011)) subtending 40° × 30° of visual angle. The design and control of task timing and visual stimulus presentation were executed with networked computers running custom written (Real-time Experimentation and Control, REX (Hays, Richmond et al. 1982)) and commercially available (Presentation, Neurobehavioral Systems) software.

### Training for Experiment 1

Monkeys were initially trained to grasp and release a touch sensitive bar to earn water rewards. After this initial shaping, a red/green color discrimination task was introduced. Red/green trials began with a bar press, and 100 ms later a small red target square (0.5°) was presented at the center of the display (over-laying a white noise background). Animals were required to continue grasping the touch bar until the color of the target square changed from red to green, this occurred randomly between 500–1,500 ms after bar touch. Rewards were delivered if the bar was released between 200–1,000 ms after the color change; releases occurring either before or after this epoch were counted as errors. All correct responses were followed by visual feedback (target square color changed to blue) after bar release and reward was delivered between 200–400 ms after visual feedback. There was a 2 second inter-trial interval (ITI), regardless of the outcome of the previous trial.

After each monkey reached criterion in the red/green task (two consecutive days with >85% correct performance) a visual categorization task was introduced. Each trial began when the animal grasped the touch bar. Next, an image (14° x 14°) appeared at the center of the screen, followed by a red cue over the center of the image. When the image presented was animate, the monkey had to release the bar before the red cue turned green to receive a liquid reward. When it was an inanimate trial, the monkey had to continue to hold the bar until the red cue turned green and then release the bar to receive a liquid reward (Figure 2). The red cue was displayed on the screen for 1-3 seconds before turning green in inanimate trials. If the monkey released during the red target and an inanimate image was presented, no reward was delivered, and the image was displayed on the screen for a 4–6 second time-out. If the monkey did not release during the inanimate image presentation within 1000 ms after the red target turned green, no reward was delivered and there was a 3 second time-out.

**Figure 2:**
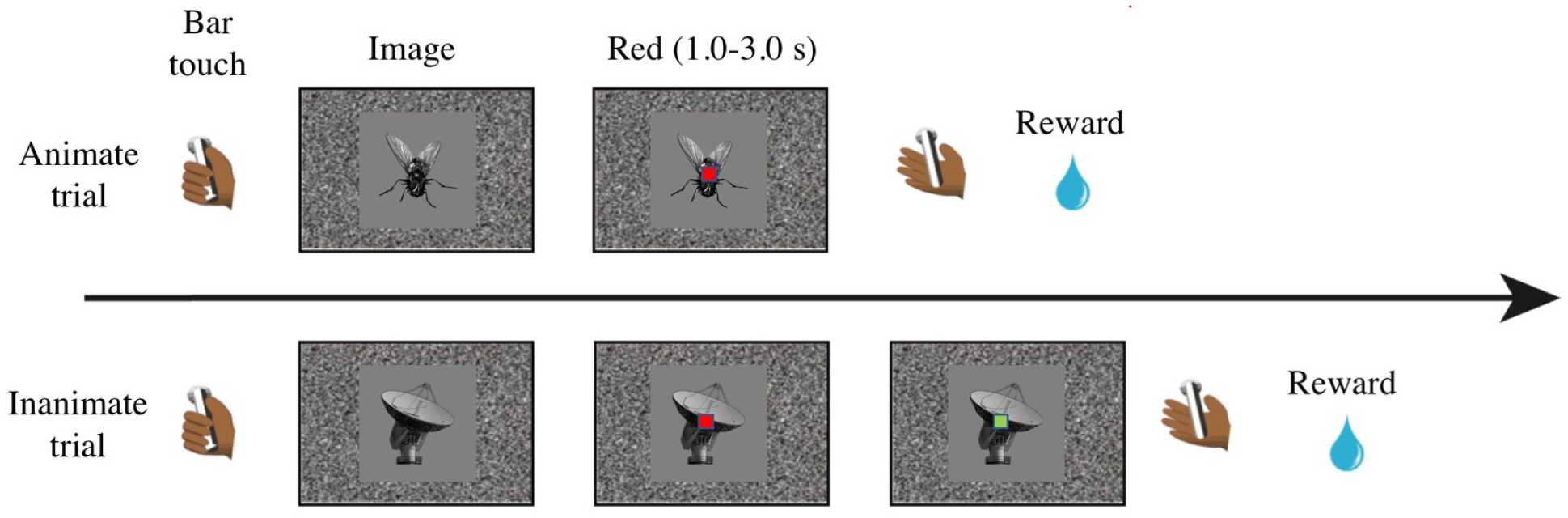
Experimental procedure. Each trial began when the animal grasped the touch bar. An image appeared at the center of the screen, followed by a red cue over the center of the image. When the image presented was animate, the monkey had to release the bar within 3 seconds of the appearance of the red cue to receive a liquid reward. When it was an inanimate trial, the monkey had to continue to hold the bar until the red cue turned green to and then release the bar to receive a liquid reward. The red cue was displayed on the screen for 1-3 seconds before turning green in inanimate trials.

If an equal drop size was used as reward for both conditions, monkeys would tend to favor a release on red because of the delay discounting effect when waiting for green. Therefore, the number of reward drops delivered for correct responses to red or green was adjusted during the training phase to reduce the bias in responding to each category for each animal. As such, the drop ratio for correct animate vs. correct inanimate trials was 1: 7 for monkey 1(M1), 1: 6 for monkey 2 (M2), and 1: 9 for monkey 3 (M3). Each monkey was trained on a repeated set of 20 animate and 20 inanimate images for several days until their choice accuracy reached above 85% accuracy for two consecutive days.

### Testing for Experiments 1 and 2

During the testing phase of Experiment 1, monkeys were tested on trial-unique sets of 100 novel animate and 100 novel inanimate intact images for 3 (M1) or 5 days (M2 and M3). After the third testing day on classifying intact images into animate and inanimate categories, M1 reached an accuracy of 91%. Due to this clear demonstration of high performance categorizing intact images, we stopped testing M1 on intact images and moved onto testing classification of synthesized images. Crucially, the training images were never shown in the testing sets, and on each testing day, monkeys were presented with a new set of unfamiliar images. Immediately after Experiment 1, monkeys were moved to Experiment 2, in which they were tested on trial-unique sets of 100 synthesized animate and 100 synthesized inanimate images (Figure 1c and 1d) for 5 days (M1, M2, M3).

### Classification analyses

The statistical significance of classification accuracy was evaluated for each monkey individually using a permutation test. For each monkey, we created a vector comprised of his responses on each trial (animate or inanimate), which we labeled as Vr, and an additional vector comprised of values representing the actual category of a trial (animate or inanimate), which we labeled as Vc. We then shuffled both the order of Vr and Vc. Then, for each row of the vectors, if the value in Vr matched that of Vc, we labeled that trial as correct and if not, as incorrect. Using this method, we calculated the overall accuracy (% correct irrespective of category), the accuracy for the animate category (% of animate trials correctly classified) and the accuracy for the inanimate category (% of inanimate trials correctly classified). The shuffling procedure was repeated 10,000 times for each monkey and for each permutation, we recorded these three accuracy values. At the end of the 10,000 permutations, each monkey had his own chance distributions (with 10,000 data points each), representing overall accuracy. Using these chance distributions, we evaluated the significance of each monkey’s actual mean classification accuracy.

### Reaction time

Since the experiments used an asymmetric design, monkeys had more time to make a decision on inanimate trials, and less time on animate trials. As such, analysis of reaction time would not yield useful information on how monkeys performed the task. Therefore, reaction time was not analyzed and presented here.

### Quantifying the amount of curvilinear and rectilinear information of the synthesized stimuli

After matching the stimuli on size, background, mean luminance and contrast, we calculated the amount of curvilinear and rectilinear information present in each image using a method presented previously in Zachariou et al. (2018) and Yue et al. (2014, 2020) (see Supplementary Materials for a detailed description).

### Logistic regression of monkeys’ performance with trial numbers

As the monkeys were rewarded when they correctly performed the categorization in the testing phase of Experiments 1 and 2, their averaged performance likely resulted from both the use of features they learned from the training images to categorize animate and inanimate images and continuous learning during the testing phase. To determine the contribution of these two factors to the overall performance, we conducted a logistic regression on each monkey’s performance using trial number as a regressor. Specifically, we regressed the monkey’s response for each trial (either right or wrong) with the trial number, in which the trial number was treated as a continuous variable. The trials in which monkeys failed to respond were excluded from the analysis. In this model, a significantly positive nonzero intercept means that the ratio of performing right over wrong is substantially larger than 1, indicating that a monkey performed the task significantly above the chance at the beginning of the experiment. A significantly larger than zero slope means their performance continuously improved as the experiment proceeded.

### Logistic regression of monkeys’ performance with curvilinear and rectilinear values of visual stimuli

To determine whether and the extent to which the amount of intermediate image features (such as curvilinear and rectilinear image features) presented in Experiments 1 and 2 contribute to monkeys’ performance, we conducted a logistic regression of monkeys’ performance (right or wrong) with the curvilinear and rectilinear values of our visual stimuli (Yue et al. 2014; Zachariou et al., 2018). The trials in which monkeys failed to respond were excluded from the analysis.

The analysis was conducted at the group level to increase the signal-to-noise ratio using MATLAB (MathWorks, Inc) with the following procedure. First, the performance from the three monkeys was concatenated to create a group response. Then curvilinear and rectilinear values for each stimulus were entered into the logistic regression model as two independent regressors. We included stimulus type (animate or inanimate) as a categorical variable in the logistic regression model to examine the interaction between amount of intermediate image features and stimulus type on monkeys’ performance. As raw responses from each monkey were used, curvilinear and rectilinear values of a stimulus that more than one monkey responded to appeared more than once in the regression model.

To determine the contribution of the amount of intermediate visual features to the monkeys’ performance, we used raw responses in a logistic regression instead of average response accuracies per stimulus in a linear regression for two reasons. First, to avoid over-estimating the influence of stimuli that only one monkey responded to, and second, to avoid creating artificially continuous responses with averaging because responses were discrete.

### Deep convolutional neural network (DCNN) training and correlation analysis

The DCNN, AlexNet (Krizhevsky et al, 2012), was imported into MATLAB, and pre-trained on the ImageNet database (Deng et al., 2009). All pre-trained weights in the first 22 layers were kept the same, while the last three layers—fully connected layer, SoftMax layer, and classification layer—were trained to classify each intact image into animate or inanimate categories. The training was conducted on the 500 intact animate and 500 intact inanimate images used in Experiment 1, using the stochastic gradient descent with momentum optimizer, minimum batch size 64, maximum epochs 20, and an initial learning rate of 10^−4^. After 300 iterations, the neural network performance converged on an accuracy of 99.9%. Then the trained neural network was tested to classify the same 1000 synthesized images used in Experiment 2 into either the animate or inanimate category.

To compute the correlation of the DCNN classification accuracies and monkeys’ response accuracies to the synthesized images in Experiment 2, we arranged the responses of the DCNN and each monkey according to the ascending order of curvilinear values of the synthesized images presented in each trial. The ordered responses were then grouped into 40 bins. The monkeys’ accuracies used for the correlation analysis were averaged across all three animals. Next, the response accuracy for each bin was calculated for the DCNN and monkeys, resulting in two sets of 40 data points. The significance of the correlation was assessed by a permutation test (10,000 iterations).

## Results

### Experiment 1: Intact images

#### 1) Overall classification accuracy for individual monkeys

During the testing phase of Experiment 1, in which novel intact images were used for the categorization task, each image was presented only once regardless of the monkeys’ responses. This eliminated the option of memorizing test images to perform the task. Across five days of testing, all monkeys performed the task significantly above chance (overall accuracy for M1: 80.88%, *p* < 0.0001; M2: 78.38%, *p* < 0.0001; M3: 76.95%, *p* < 0.0001). The statistical significance was determined by the permutation test (see Methods). The overall response rate was 99.64% for M1, 73.43% for M2, and 98.86% for M3.

Upon closer inspection of the data we found that M2 memorized all 40 training images to perform the categorization task. Thus, in the first day of testing, M2 was learning the categorization task. After eliminating data from this day, overall performance was 85.64% (*p* < 0.001), and overall response rate was 73.3%. Unless stated otherwise, subsequent analyses used M2’s testing data from day 2 to day 5 only. Data from all five days of testing are included in Supplementary Figure 1.

The data show that monkeys were able to successfully classify intact images that they had no previous experience with into animate and inanimate object categories, suggesting that image-based features distinguishing the two categories played a significant role in monkeys’ categorization performance.

#### 2) Generalization and learning effect for individual monkeys

Because monkeys were given a liquid reward whenever they categorized images correctly in the testing phase, their overall performance could have resulted from continuously learning to categorize testing images as animate and inanimate due to reward feedback. In other words, significantly above-chance performance in the testing phase may not have captured the full picture of the monkeys’ complex performance processing. Their performance could have more to do with this continuous feedback than with generalizing visual features learned during the training set to categorize the testing images. To separate the effect of generalization from the effect of learning during the testing phase, we performed a logistic regression (see Methods) on a single-trial basis to quantify the generalization as the intercept and learning as the slope of the regression model. We anticipated that, if there were a generalization effect, then the intercept of the logistic regression model would be significantly greater than zero, and if there were a learning effect, then the slope of the regression model would be significantly greater than zero.

Monkeys were able to use the information they learned during training to perform the categorization task on unfamiliar images at the onset of the testing phase, as shown in Table 1, where the intercept of the logistic regression is shown to be significantly above chance for all three monkeys. The slope of the logistic regression was positive and significantly different from zero in all monkeys, indicating that performance improved as testing progressed. All three monkeys’ performance was significantly predicated by trial number, as shown in Figure 3 and Table 1 (for M1: χ^2^ (595) = 58.545, *p* = 1.98 × 10^−14^; M2: χ^2^(584) = 18.361, *p* = 1.828 × 10^−5^; M3: χ^2^(986) = 13.252, *p* = 2.72 × 10^−4^), further indicating that monkeys continued to learn during the testing phase, improving their performance even though each image was presented only once.

**Table 1.** Logistic regression results from Experiment 1.

| Monkeys | Logistic regression |  |
| --- | --- | --- |
|  | Intercept | Slope |
| M1 | 0.359 ($p = 2.4 \times 10^{-2}$) | $2.951 \times 10^{-2}$ ($p = 5.241 \times 10^{-13}$) |
| M2 | 1.086 ($p = 2.921 \times 10^{-9}$) | $1.94 \times 10^{-2}$ ($p = 2.746 \times 10^{-5}$) |
| M3 | 0.809 ($p = 1.211 \times 10^{-10}$) | $6.368 \times 10^{-3}$ ($p = 2.969 \times 10^{-3}$) |

**Figure 3.**
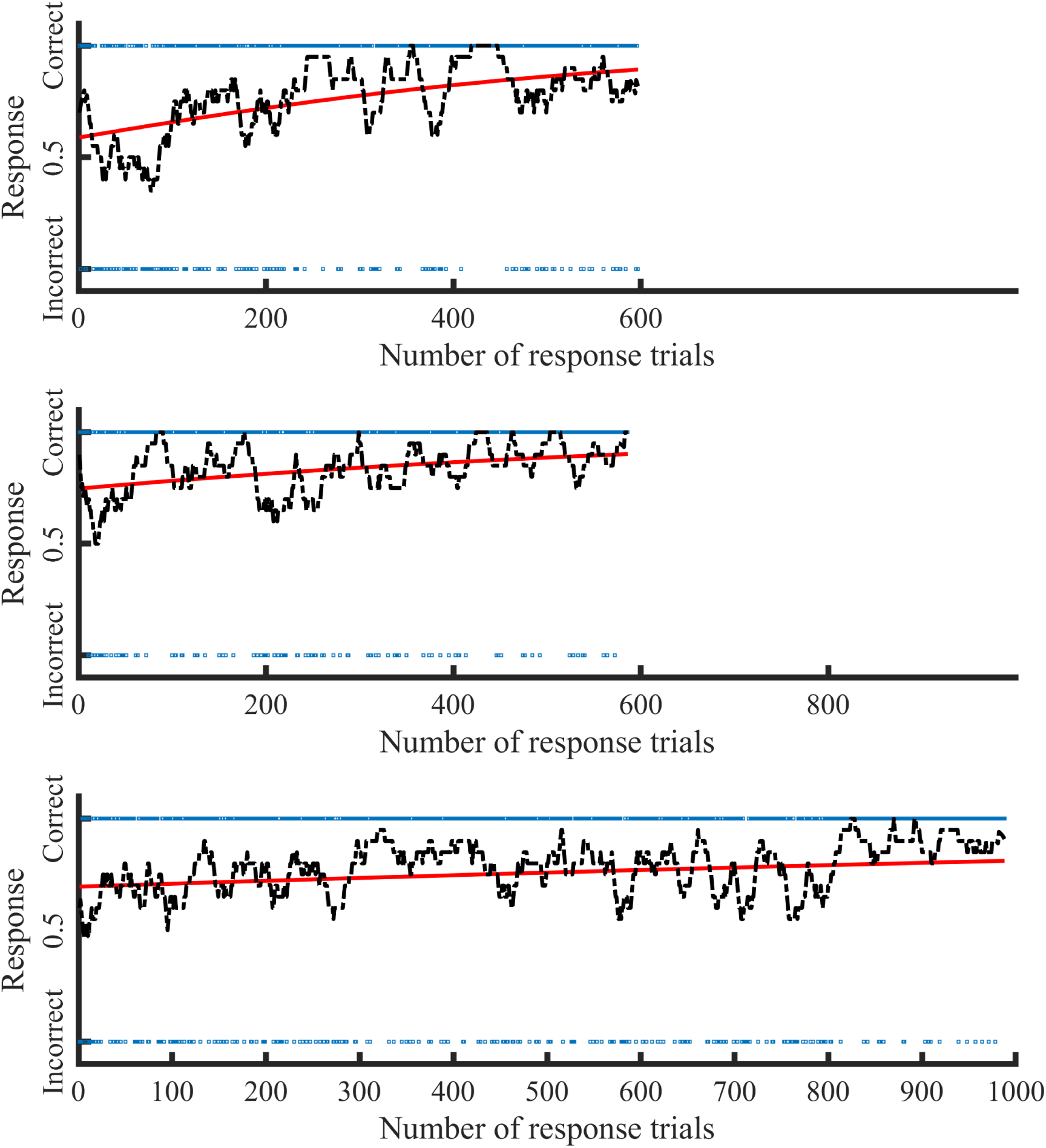
The logistic regression results of Experiment 1 for M1 (top), M2 (middle), and M3 (bottom). The x-axis represents the number of response trials (trials without responses were removed), and the y-axis represents the monkey’s response. The monkeys’ responses for each trial are shown as blue dots, which appears as a blue line because of the large number of trials. The red line represents the predicted response probability produced from the logistic regression analysis. The black dotted line represents the response accuracy of a moving average of 20 trials, which is for illustration purposes only and not used for calculating logistic regression. The intercepts of the regression lines for all three monkeys are larger than 0.5, indicating that all three monkeys were able to generalize from the training set to the testing set. The regression line increased along with the trial numbers, suggesting that monkeys continued to learn during the testing phase to improve their performance. M1 was tested only for three days; therefore, it has only 600 trials. M2 was tested for five days, but data from the first day were removed from the logistic regression.

Taken together, the significantly above-chance performance and significant generalization effect in categorizing the intact novel images suggests that all three monkeys learned to distinguish between the two categories during the training phase (M1 and M3) or after the first day of testing (M2), by generalizing the features learned from the small set of training images to the unfamiliar images in the larger testing set.

#### 3) Contribution of curvilinear and rectilinear features to monkeys’ performance at the group level

We aimed to understand the extent to which the amount of intermediate image features, specifically curvilinear and rectilinear features (see Methods), present in the images in Experiment 1 contributed to the monkeys’ performance on the categorization task. To answer this question, we conducted a logistic regression analysis of curvilinear and rectilinear values with monkeys’ performance, which was performed at the group level to increase the signal-to-noise ratio (see Methods).

We found that the amount of intermediate image features in the intact images significantly predicted monkeys’ performance (main effect: χ^2^ (2768) = 107.4, *p* = 1.450 × 10^−21^), suggesting that the amount of intermediate image features might assist them in categorizing intact images into animate and inanimate groups. Furthermore, we found that curvilinear values of intact images significantly predicted monkeys’ performance (beta = 0.974, *p* = 0.031), but rectilinear values did not (beta = −0.4817, *p* = 0.272). There was a significant interaction between the curvilinear values and the stimulus category (beta = −2.21, *p* = 1.118 × 10^−4^), indicating that curvilinear values predicted monkeys’ performance in animate trials differently than on inanimate trials. Figure 4 shows the functional relationship between curvilinear values and monkeys’ performance across animate and inanimate trials, which was produced from the logistic regression model. As the amount of curvilinear information in an image increased, monkeys’ performance increased for animate images and decreased for inanimate images.

**Figure 4.**
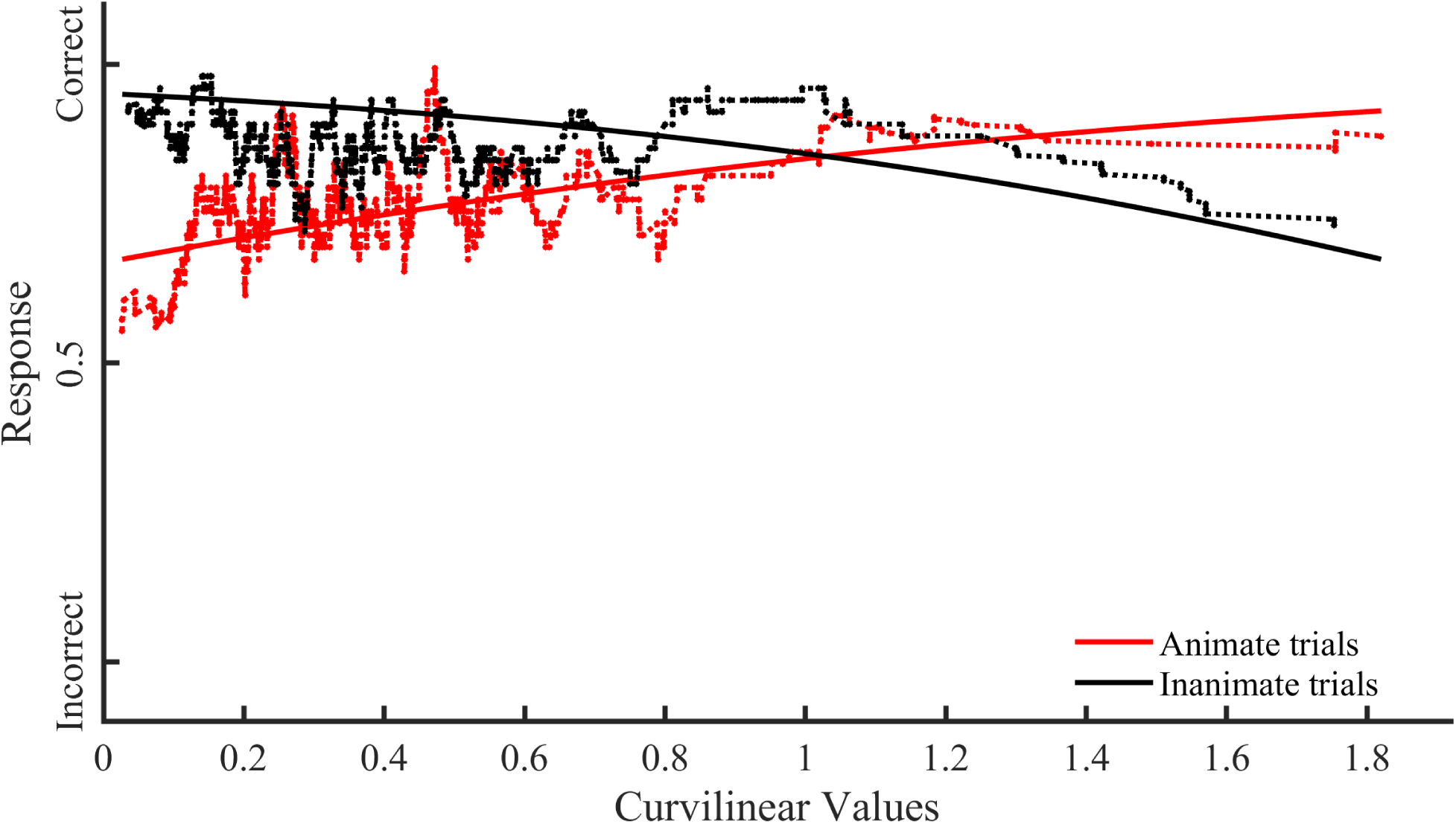
Functional relationship between the amount of curvilinear information present in visual stimuli and monkeys’ performance across stimulus category in Experiment 1. The x-axis represents the curvilinear values of the stimuli. The y-axis represents the response probability of the monkeys’ performance. The solid lines represent the response probability to visual stimuli calculated with the logistic regression model that was created using the monkeys’ group raw response. The dotted lines represent a moving average of 60 trials, which is for illustration purposes only and was not used for fitting the logistic regression model. The red line represents the response probability resulting from the logistics regression fitting for the animate trials. The black line represents the response probability resulting from the logistics regression fitting for the inanimate trials

**Figure 5:**
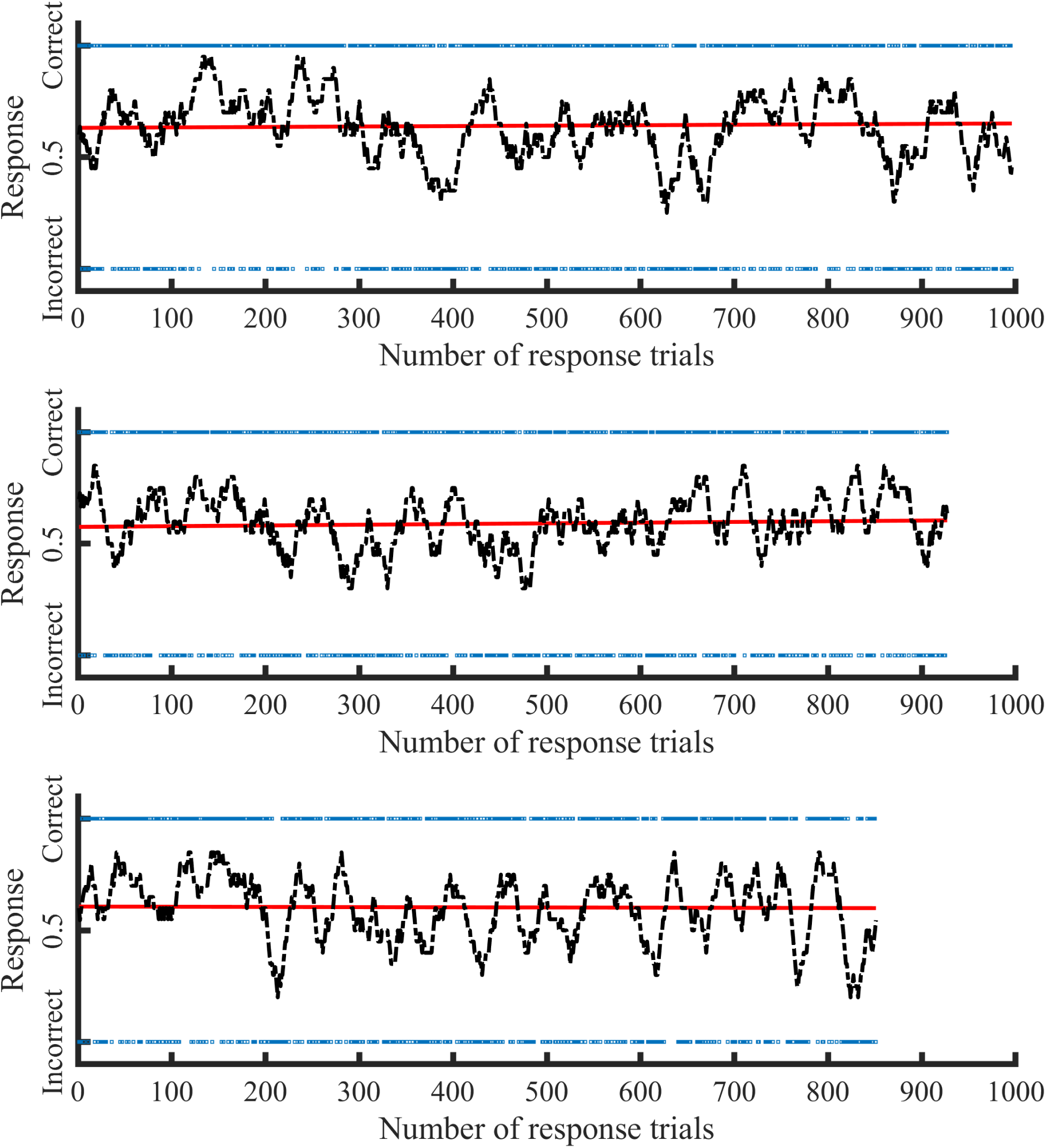
The logistic regression results of Experiment 2 for M1 (top), M2 (middle), and M3 (bottom). Axes are the same as those used in Figure 3. As shown in Table 2, all three monkeys showed significant generalization but no learning effects. These results suggest that the monkeys employed some image features distinguishing intact animate images from intact inanimate images to categorize the synthesized images as animate or inanimate.

These results suggest that, in addition to recognizing local or global features that the monkeys had learned during daily training, monkeys may also have used the amount of curvilinear image features present in the stimuli to categorize objects into animate and inanimate groups.

### Experiment 2: Synthesized images

#### 1) Overall classification accuracy for individual monkeys

The monkeys were never trained to categorize the synthesized images presented in Experiment 2. Furthermore, the synthesized images were each shown only once, regardless of the monkeys’ responses. As shown in Figure 3B, all three monkeys performed the categorization task significantly above chance (overall accuracy for M1, 64.48%, *p* < 0.0001; M2, 59.10%, *p* < 0.0001; M3, 60.27%, *p* < 0.0001). The overall response rate was 99.6% for M1, 92.7% for M2, and 85.1% for M3. Although the overall classification accuracies were lower than those for the intact images in Experiment 1, the significant above-chance performances suggest that the image features distinguishing the two groups of synthesized images provided sufficient information for monkeys to classify the images into the two categories.

#### 2) Generalization and learning effect for individual monkeys

To provide a parallel analysis to the one performed in Experiment 1, we ran a logistic regression to evaluate if the monkeys’ overall accuracies for categorizing the synthesized images resulting from generalizing visual features learned from the intact images to the synthesized images and/or continuous learning. We found that the intercept, but not the slope, of the logistic regression model was significant for all three monkeys, as shown in Table 2. Performance was not significantly determined by test trial number for any monkeys (for M1: χ^2^ (994) = 0.365, *p* = 0.546; M2: χ^2^(925) = 0.340, *p* = 0.560; M3: χ^2^(849) = 0.032, *p* = 0.859), indicating that monkeys’ performance did not improve as testing progressed. These results reveal that, at the onset of Experiment 2, all three monkeys used information they learned on the categorization task in Experiment 1 to classify the synthesized images as animate and inanimate objects.

**Table 2.** Logistic regression result of Experiment 2.

| Monkeys | Logistic regression |  |
| --- | --- | --- |
|  | intercept | Slope |
| M1 | 0.533 ($p = 2.038 \times 10^{-6}$) | $9.095 \times 10^{-5}$ ($p = 0.521$) |
| M2 | 0.313 ($p = 1.480 \times 10^{-2}$) | $1.150 \times 10^{-3}$ ($p = 0.606$) |
| M3 | 0.428 ($p = 4.816 \times 10^{-3}$) | $-1.930 \times 10^{-5}$ ($p = 0.919$) |

#### 3) Contribution of curvilinear and rectilinear features to monkeys’ performance at the group level

To examine the extent to which the amount of intermediate visual features contributed to monkeys’ performance in Experiment 2, we used the same testing procedure as Experiment 1 but with synthesized images.

We found a significant main effect of the amount of curvilinear and rectilinear image features on monkeys’ performance (χ^2^ (2768) = 177.160, *p* = 2.160 × 10^−36^). Furthermore, both curvilinear and rectilinear values of synthesized images significantly predicted monkeys’ performance (curvilinear: beta = 1.617, *p* = 2.615 × 10^−7^; rectilinear: beta = −1.257, *p* = 5.865 × 10^−4^). However, the data suggested that the amount of curvilinear image features present in the synthesized images played a more dominant role than the amount of rectilinear image features. To test this hypothesis, we performed a regression Wald test to examine whether the curvilinear coefficient was significantly different from the rectilinear coefficient. The curvilinear coefficient was significantly larger than the rectilinear coefficient (Wald test: χ^2^(1) = 19.938, *p* = 7.994 × 10^−6^), indicating that the amount of curvilinear image features present in the synthesized images was more informative for the categorization task than the amount of rectilinear image features. As such, the following analysis of interaction between the amount of intermediate image features and stimulus category was focused on the contribution of the amount of curvilinear image features on monkeys’ performance across stimulus categories. Results of the analysis of the interaction effect between the amount of rectilinear image features with stimulus category are shown in Supplementary Figure 2.

We observed a significant interaction between the curvilinear values of stimuli and stimulus category (beta = −4.040, *p* = 1.672 × 10^−20^). Monkeys’ performance on synthesized images increased when curvilinear values increased in the animate trials but decreased in the inanimate trials (Figure 6); similar to what we observed in Experiment 1 (Figure 4). These data indicate that the more curvilinear information present in an animate image, the more likely it was to be categorized correctly, whereas the opposite is true for inanimate images.

**Figure 6.**
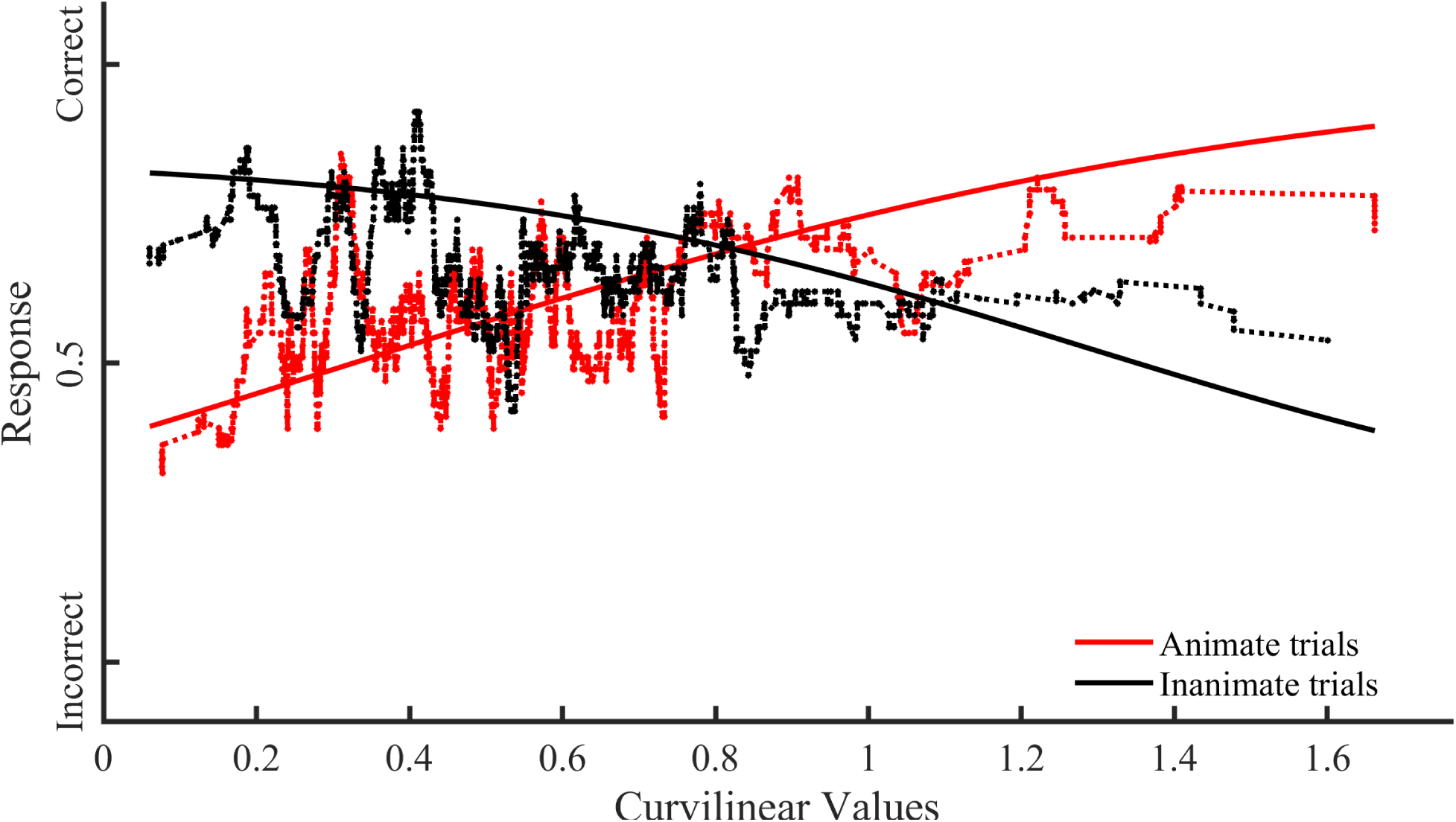
Functional relationship between amount of curvilinear information present in the visual stimuli and monkey’s group performance across stimulus category in Experiment 2. The x-axis represents the curvilinear values of visual stimuli. The y-axis represents the response probability of the monkeys’ performance. The solid lines represent the response probability to visual stimuli calculated with the logistic regression model that was created using the monkeys’ group raw response. The dotted lines represent a moving average of 60 trials, which is for illustration purposes only. The red line represents the response probability resulting from the logistics regression fitting for the animate trials. The black line represents the response probability resulting from the logistics regression fitting for the inanimate trials.

#### 4) Correlation of monkeys’ performance with DCNN performance at the group level

Because monkeys were never trained to classify synthesized images into animate and inanimate categories, the possibility remained that monkeys categorized the images into two groups using differences between synthesized images that were entirely unrelated to the animate and inanimate category but happened to coincide with the two categories in the set of testing images used. As such, we used the DCNN to address this concern (see Methods). The network was trained to classify the 1000 intact images used in Experiment 1 into animate and inanimate categories and then tested on the categorization task with the 1000 synthesized images used in Experiment 2 (see Methods). We found a significant positive correlation of the DCNN’s categorization performance with the monkeys’ group performance (r = 0.739, *p* = 5.0502 × 10^−8^) (Figure 7), suggesting that the monkeys performed the animate vs. inanimate categorization in Experiment 2, when the global form in the images was distorted beyond recognition. These data provided further evidence that the monkeys used image features distinguishing intact animate and inanimate images to categorize the synthesized images.

**Figure 7:**
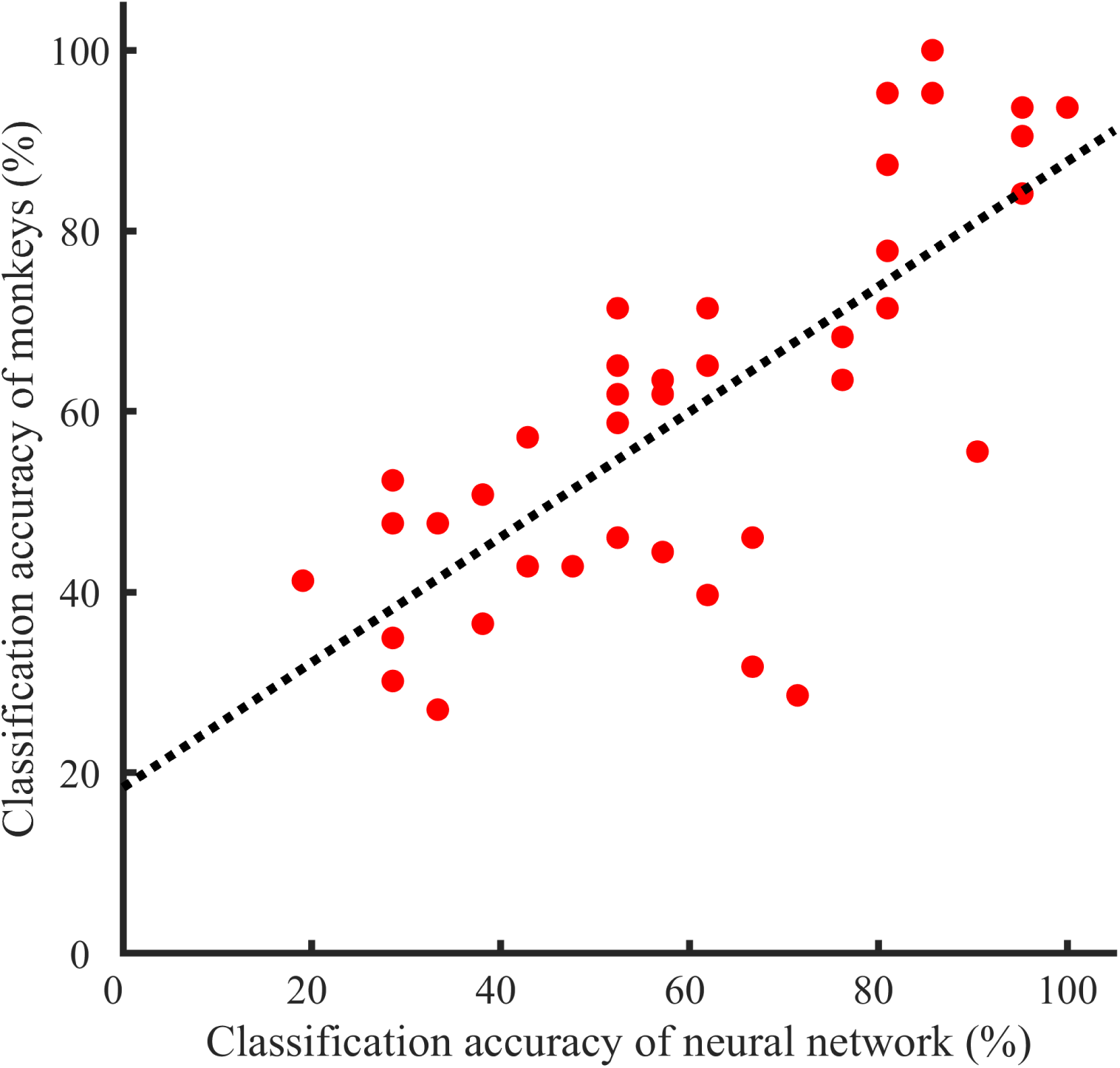
Correlation of monkeys’ response accuracies with DCNN classification accuracies. To compute the correlation of the DCNN classification accuracies and monkeys’ response accuracies to the synthesized images, we arranged the responses of the DCNN and each monkey according to the ascending order of curvilinear values of the synthesized images. The monkeys’ accuracies used for the correlation analysis were averaged across all three animals. The ordered responses were then grouped into 40 bins. Next, the response accuracy for each bin was calculated for the DCNN and monkeys separately, resulting in two sets of 40 data points. Each red dot represents the classification accuracy for each bin. We observed a significant correlation between monkeys’ response accuracies and DCNN classification accuracies (r = 0.739, *p* = 5.0502 × 10^−8^), indicating that monkeys performed the animate vs. inanimate categorization.

## Discussion

This study investigated the contributions of both training and image-based features to the perceptual categorization of animacy. In Experiment 1, we found that naïve monkeys trained to categorize a small set of animate and inanimate images classified a large set of unfamiliar images into animate and inanimate categories with high accuracy. In Experiment 2, we tested whether image-based features that differ between the two object categories in the statistics of natural environments, i.e. curvilinear and rectilinear information (Kurbat, 1997; Levin et al. 2001; Perrinet and Bednar, 2015; Long et al. 2017; Zachariou et al., 2018), determined the monkeys’ classification accuracy. We created sets of synthetic animate and inanimate images using an algorithm that significantly distorted the global shape of the original images while maintaining the original images’ intermediate features (Portilla and Simoncelli, 2000). The monkeys’ classification accuracy on these synthesized images was still significantly above chance and correlated with the amount of curvilinear information present in the stimuli. These data indicate that image-based features, in this case curvilinearity, can be used to distinguish animate from inanimate objects in the absence of global shape information without prior training.

As monkeys raised in the laboratory have limited experiences with objects that humans are otherwise familiar with, they are ideal candidates to study the contribution of experiences and image-based features to the emergence of perceptual categorization (e.g. Arcaro & Livingstone, 2017). Our results show that monkeys performed an animacy categorization task with intact images significantly above chance at the very beginning of the test phase of Experiment 1, suggesting that monkeys used what they had learned during training to classify novel images of objects, with which they had no previous experience, into animate and inanimate categories. Further, the curvilinear values of intact images had a significant interaction with stimulus category, and significantly predicted the monkeys’ performance. These findings indicate that image-based features that are predictive of each category provide substantial information that monkeys can use to distinguish the two categories with little training. In other words, experience interacting with objects may not be the only origin of behavioral categorization of animacy in monkeys.

To confirm this, using the synthesized images in Experiment 2, we eliminated local features (faces, ears, etc.) that monkeys might have been familiar with and could have used to classify the images into animate and inanimate categories. We found that the monkeys were able to perform the categorization of the synthesized images significantly above chance, which indicates that the image-based features were sufficient for the emergence of perceptual categorization. It is worth noting that human participants also classified synthesized images similar to those used in this experiment into animate and inanimate categories with significant above-chance accuracy (Zachariou, et al., 2018; Long et al., 2017). Although humans and monkeys do not share the collective experience of what and how objects are encountered in daily life, they perform similarly when classifying synthesized images into animate and inanimate categories (Figure 6, Figure 3 in Zachariou, et al., 2018), which suggests that image-based feature differences could play a critical role in the emergence of perceptual categorization abilities across species. Together, our findings provide strong evidence in support of the hypothesis that perceptual categorization can emerge from image-based features that are predictive of each category in the natural statistics of the visual environment.

Recent fMRI studies (Long et al., 2018; Yue et al, 2020) have shown that visual cortical areas selective for curvilinear features encompass animate-processing visual areas while those selective for rectilinear features encompass inanimate-processing visual areas. These results provide neural evidence to support the current finding that the processing of image-based features, such as curvilinearity, interacts with the representation of animate and inanimate categories.

Overall, monkeys categorized the intact object images with significantly greater accuracy than the synthesized images. However, for synthesized images with high curvilinear values (in the range of 1.4 – 1.6), monkeys’ classification accuracy for the animate category could reach above 80% which is comparable to the classification accuracy for intact images (Figure 6). This illustrates that monkeys could achieve high accuracy when synthesized images with extreme curvilinear values were used as stimuli. Thus, the overall difference in classification accuracy between the intact and synthesized images does not argue against the idea that image-based features play a significant role in determining perceptual categorization.

The primate visual system takes significant time to fully mature postnatally (Gilmore et al., 2018; Ellemberg et al., 1999; Kovacs et al., 1999). During development, young infants view the world as consisting not of coherent objects but instead visual pieces that move in unpredicted ways (Hyvärinen, et al., 2014). In such a fragmented visual world, differentiating animate from inanimate objects would be challenging. Infants who can differentiate animate from inanimate objects would have a better chance to avoid being harmed by animals to survive than those who cannot. Through natural selection, our brains may have evolved the capacity to differentiate animate and inanimate objects quite quickly, first based on sensory information that represents visual statistics of the natural environment. Experience with objects would play a significant role in later life to further differentiate categories. Our data provide evidence to support this hypothesis by showing that monkeys (as well as humans (Zachariou, et al., 2018)) are able to classify synthesized images that: 1) neither species has experience with; and 2) have similar statistics as the natural original images, into animate and inanimate categories significantly above chance by using the degree of curvilinearity in the images. This hypothesis raises many interesting questions. For which object categories and with which image features is the primate brain biased to use image-based differences for perceptual categorization, and under what conditions? The answers to such questions are critical to understand the functional and anatomical organization of the primate visual system.

## Acknowledgments

This work was supported by the NIMH Intramural Research Program.

## Supplmentary Materials

**Figure 1.**
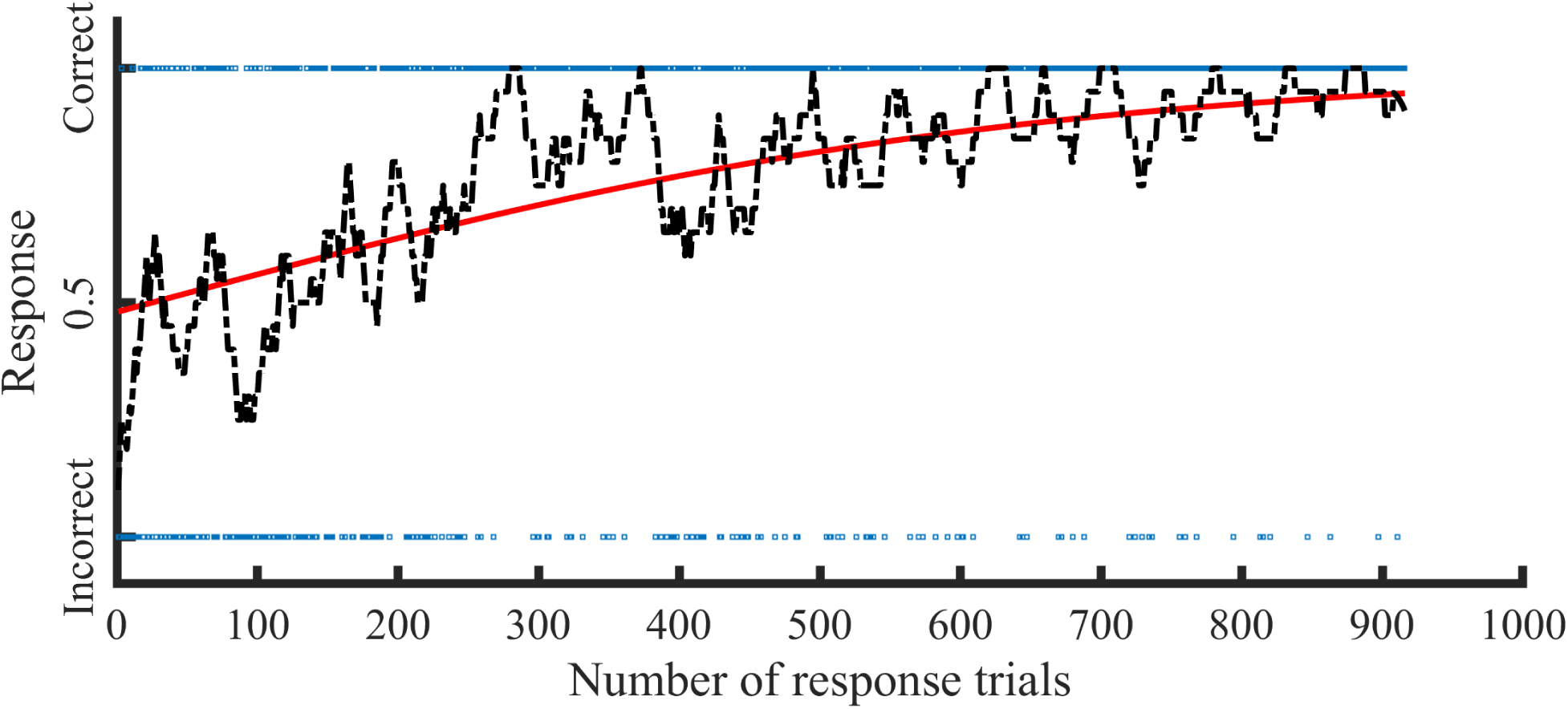
The logistic regression results of the experiment 1 for M2. The x-axis represents the number of response trials for all five days, and the y-axis represents the monkey’s response. The monkey responses for each trial are shown as blue dots, which appears as a blue line because of the large number of trials. The red line represents the predicted response probability produced from the logistic regression analysis. The black dot line represents the response accuracy of a moving average of 20 trials, which is for illustration purposes only and not used for calculating logistic regression.

**Figure 2.**
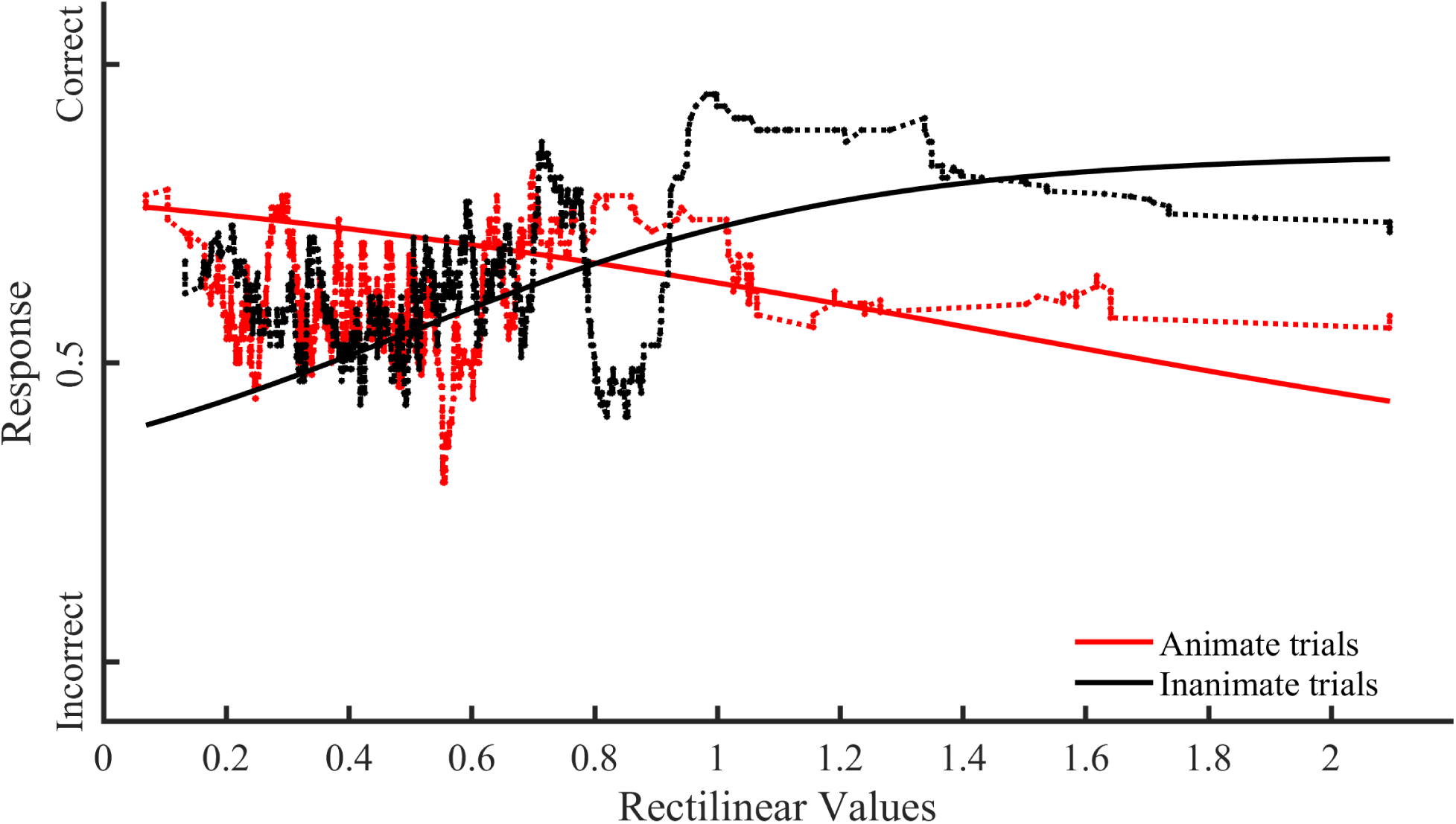
Functional relationship between amount of rectilinear information present in the visual stimuli and monkey’s group performance across stimulus category in Experiment 2. The x-axis represents the rectilinear values of visual stimuli. The y-axis represents the response probability of the monkeys’ performance. The solid lines represent the response probability to visual stimuli calculated with the logistic regression model that was created using the monkeys’ group raw response. The dotted lines represent a moving average of 60 trials, which is for illustration purposes only.

